# Therapy-associated mutagenesis at CTCF binding sites is shaped by chromatin context and DNA repair capacity

**DOI:** 10.64898/2026.04.15.718780

**Authors:** Kevin C.L. Cheng, Zoe P. Klein, Jigyansa Mishra, Alexander T. Bahcheli, Benjamin H. Lok, Trevor J. Pugh, Jüri Reimand

## Abstract

Genotoxic cancer therapies introduce DNA damage that can be fixed as somatic mutations in surviving tumor cells. However, the impact of therapy-associated mutagenesis on regulatory elements remains unclear. CTCF binding sites (CBS) are chromatin architectural elements that exhibit recurrent localized mutation enrichment in cancer genomes. We asked whether treatment exposure is associated with increased mutagenesis at CBS in 4,870 whole-genome sequences from metastatic tumors across 17 cancer types and 45 therapies. Radiotherapy and trifluridine exposure in metastatic colorectal cancer were associated with increased mutation enrichment at CBS. This enrichment was pronounced at motif-containing sites and in low-expression or late-replicating genomic contexts. Alterations in DNA damage response genes, including BRCA2, were associated with increased CBS mutation enrichment following radiotherapy. Together, these findings indicate that therapy-associated mutagenesis at CTCF binding sites is shaped by chromatin context and DNA repair capacity, extending the mutational consequences of cancer treatment to regulatory genome architecture.

## Background

Many cancer therapies exert cytotoxic effects by inducing DNA damage [1,2]. In surviving tumor cells, imperfect repair of therapy-induced lesions can result in fixed somatic mutations that leave detectable mutational footprints in the cancer genome [3]. However, mutation accumulation across the cancer genome is non-uniform [4,5] and is shaped by replication timing, chromatin organization, transcriptional activity, transcription factor binding, and local sequence context [6–13]. At fine genomic scales, protein-DNA interactions and local repair accessibility can influence whether damage is successfully repaired or gives rise to mutations [14]. Understanding how therapy-induced DNA damage interacts with these layers of genomic organization is essential for interpreting the tumor evolutionary consequences of cancer treatment.

CTCF binding sites are regulatory elements in non-coding DNA that exhibit recurrent localized mutation enrichment across multiple cancer types [10,15–19]. CTCF is a key architectural protein that organizes chromatin domains and topologically associated domains (TADs), and its binding sites anchor regulatory interactions across the genome [20]. Our previous work demonstrated that constitutively active CTCF binding sites, defined as sites bound by CTCF across diverse tissue types, exhibit the strongest localized mutation enrichment in multiple cancer types [19]. Notably, transcription factor binding at CTCF binding sites has been shown to impair nucleotide excision repair, suggesting that sustained CTCF occupancy may constrain local repair and increase susceptibility to mutation [14,17]. In addition, we previously observed that highly mutated binding sites of CTCF are frequently co-bound by TOP2B, a topoisomerase involved in the generation and repair of DNA double-strand breaks and a known off-target of certain chemotherapeutic agents [21]. Because these sites contribute to higher-order chromatin architecture, mutations at CTCF binding sites have the potential to disrupt regulatory domains and affect multiple genes [10,22,23]. Mutational consequences of several chemotherapeutic agents and radiotherapy have been characterized in cancer genomes [3,24–27]. However, it remains unclear how therapy-associated mutagenesis is distributed across specific classes of regulatory elements and whether DNA damage is preferentially fixed at sites with constrained repair accessibility. In particular, the impact of genotoxic therapies on non-coding regulatory elements, including CTCF binding sites, has not been systematically evaluated in large cohorts of treated tumors.

Given the susceptibility of CTCF binding sites to somatic mutagenesis, we hypothesized that genotoxic cancer therapies may preferentially increase mutation enrichment at these sites in surviving tumor cells. To address this question, we analyzed whole-genome sequencing data from a large cohort of metastatic tumors spanning multiple major cancer types, systematically assessing therapy-associated mutation enrichment at constitutively active CTCF binding sites. We identified associations between genotoxic therapies and increased mutation enrichment at CTCF binding sites in metastatic colorectal cancer, and found that these effects were modulated by local chromatin context and tumor-specific alterations in DNA damage response pathways. Together, these findings suggest that cancer therapies can reshape mutagenesis at regulatory elements in a manner influenced by both genomic architecture and repair capacity.

## Results

### Cancer therapies are associated with localized mutation enrichment at constitutively active CTCF binding sites

We analyzed whole-genome sequencing (WGS) data for 4870 metastatic tumors across 17 cancer types from the Hartwig Medical Foundation (HMF) cohort [28] (**Table S1**). To assess whether cancer therapies are associated with localized mutagenesis at CTCF binding sites (CBS), we focused on approximately 30,000 constitutively active binding sites of CTCF identified across many human tissue types [19] (**Table S2**). This subset was defined as CTCF-bound in a majority of cell types profiled in the ENCODE project [29] and was selected for our analysis due to a strong mutation enrichment in cancer genomes [19,21]. CBS were defined as central sequences of 50 basepairs around ChIP-seq peak midpoints, and somatic mutation counts within these regions were compared to matched flanking sequences as local background controls. Mutation counts were modeled using RM2, a negative binomial model that accounts for trinucleotide sequence context and megabase-scale background mutation burden, to quantify CBS-specific mutation enrichment or depletion [19].

We first performed a systematic computational analysis to identify pairs of cancer types and therapies associated with localized changes in mutation enrichment at CBS (**Figure 1a, Figure S1**). We evaluated 45 individual therapies and 15 therapy groups across 17 cancer types, requiring a minimum of 25 treated and 25 untreated tumors in each pair to ensure statistical power (**Tables S1, S3**). For each eligible cancer-therapy pair, we used RM2 to quantify mutation enrichment at CBS relative to immediately flanking regions, which served as a matched local background. We then tested statistically whether treatment exposure was associated with a change in the CBS-to-flank mutation enrichment ratio, followed by multiple-testing adjustment of significance estimates.

**Figure 1.**
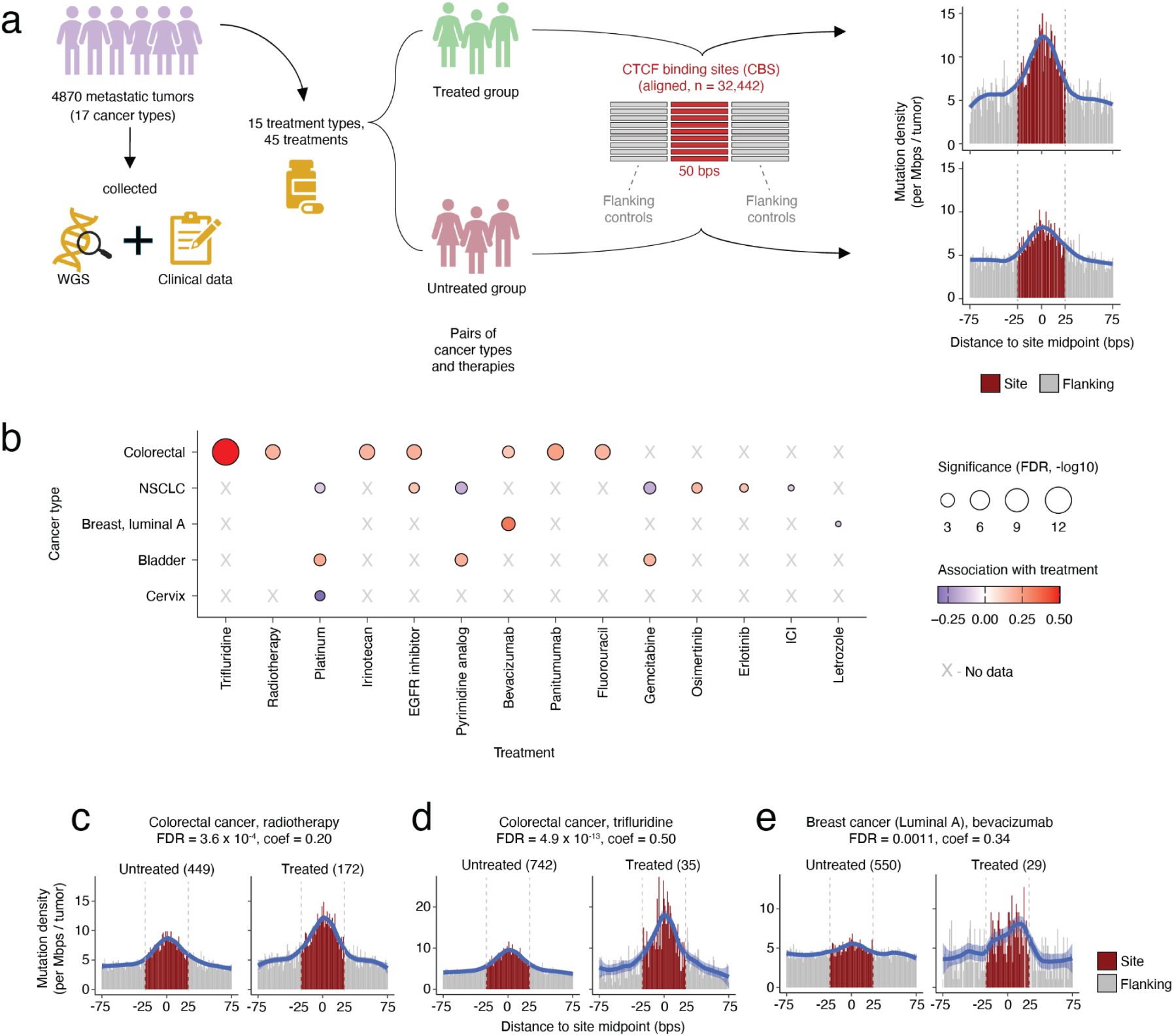
Systematic analysis identifies cancer therapies associated with localized mutation enrichment at constitutive CTCF binding sites (CBS) in metastatic cancers. **(a)** Overview of the analysis of cancer therapies and CBS mutation enrichment. **(b)** Significant associations of therapies and cancer types identified in the systematic analysis. Each dot represents a pair of cancer type and therapy, tested for differences in CBS mutation enrichment between treated and untreated tumors. Dot color indicates the direction and magnitude of the treatment-by-site interaction coefficient, where positive values indicate increased CBS mutation enrichment in treated tumors. Only pairs of cancer types and therapies meeting inclusion criteria are shown (at least 25 tumors in both treated and untreated groups). Crosses indicate insufficient sample size for testing. Mutation enrichment at CBS relative to flanking regions was modeled using the RM2 framework with negative binomial regression, while accounting for trinucleotide sequence context and 1 Mb background mutation burden as covariates, and adjusted for multiple testing (chi-square test, FDR < 0.05). **(c-e)** Representative examples of therapy-associated CBS mutation enrichment. Mutation density was aggregated by aligning CBS at their midpoints. The core CBS regions (±25 bp or 50 bp in total; shown in dark red) were compared to flanking regions (50 bp upstream and downstream; shown in grey). Blue lines show LOESS-smoothed mutation density (span = 0.4); blue shading indicates standard error of LOESS fit. Acronyms: false discovery rate (FDR); non-small cell lung cancer (NSCLC); immune checkpoint inhibition (ICI).

Across this systematic analysis, we identified 20 significant associations involving 13 therapies and five cancer types (FDR < 0.05; **Figure 1b, Table S1**). Among these, metastatic colorectal cancer exhibited multiple strong and statistically robust associations, with several therapies showing significant increases in mutation enrichment in CBS relative to flanking genomic regions. In metastatic colorectal cancer, radiotherapy and trifluridine exposure showed the strongest associations with CBS mutation enrichment. Radiotherapy-treated colorectal tumors (n = 172) showed 1.33-fold mutation enrichment at CBS relative to flanking genomic sequences, compared to 1.24-fold enrichment in 449 untreated tumors (FDR = 3.6 x 10⁻⁴ from RM2; **Figure 1c**). Trifluridine exposure was associated with a 1.52-fold CBS mutation enrichment increase in 35 treated tumors, compared to 1.22-fold increase in 742 untreated tumors (FDR = 4.9 x 10^-13^, **Figure 1d**). Several additional therapies used commonly to treat patients with colorectal cancer, such as irinotecan, fluorouracil, bevacizumab, and anti-EGFR agents, also showed significant associations with CBS mutation enrichment (**Figure 1b, Table S1**). However, because these therapies were frequently administered in combination, their individual contributions require further deconvolution. These results indicate that constitutively active CBS are a prominent target of therapy-associated mutagenesis in metastatic colorectal cancer.

Beyond colorectal cancer, we identified a few significant therapy associations with CBS mutation enrichment in other cancer types. For example, bevacizumab exposure was associated with increased CBS mutation enrichment in luminal A breast cancer (FDR = 0.0011; **Figure 1e**). However, these associations generally showed modest effect sizes or were supported by limited sample numbers, reducing their suitability for detailed downstream analyses (**Table S1**). In contrast, radiotherapy and trifluridine exposure in metastatic colorectal cancer were supported by larger cohorts (**Table S4**) and showed strong and reproducible CBS mutation enrichment, allowing rigorous assessment of independent treatment effects. Therefore, we focused our next analyses on treatment associations in metastatic colorectal cancer to investigate factors shaping therapy-associated CBS mutagenesis.

### Radiotherapy and trifluridine associate with mutation enrichment at CTCF binding sites in colorectal cancer

Because many patients with colorectal cancer received multiple therapies, there was substantial overlap among treatments showing significant associations in the initial systematic analysis. To disentangle their independent contributions, we fitted a joint model including the six colorectal cancer therapies identified from the initial analysis (trifluridine, radiotherapy, irinotecan, bevacizumab, fluorouracil, and anti-EGFR therapy). In this model, radiotherapy and trifluridine remained significantly associated with increased CBS mutation enrichment (*P* = 0.0050 and *P* = 0.036, respectively, **Figure 2a**), whereas the other therapies were largely attenuated and no longer significant. These results indicate that the associations of radiotherapy or trifluridine with CBS mutation enrichment are not explained by co-treatment with other agents. We therefore focused subsequent analyses on these two therapies.

**Figure 2.**
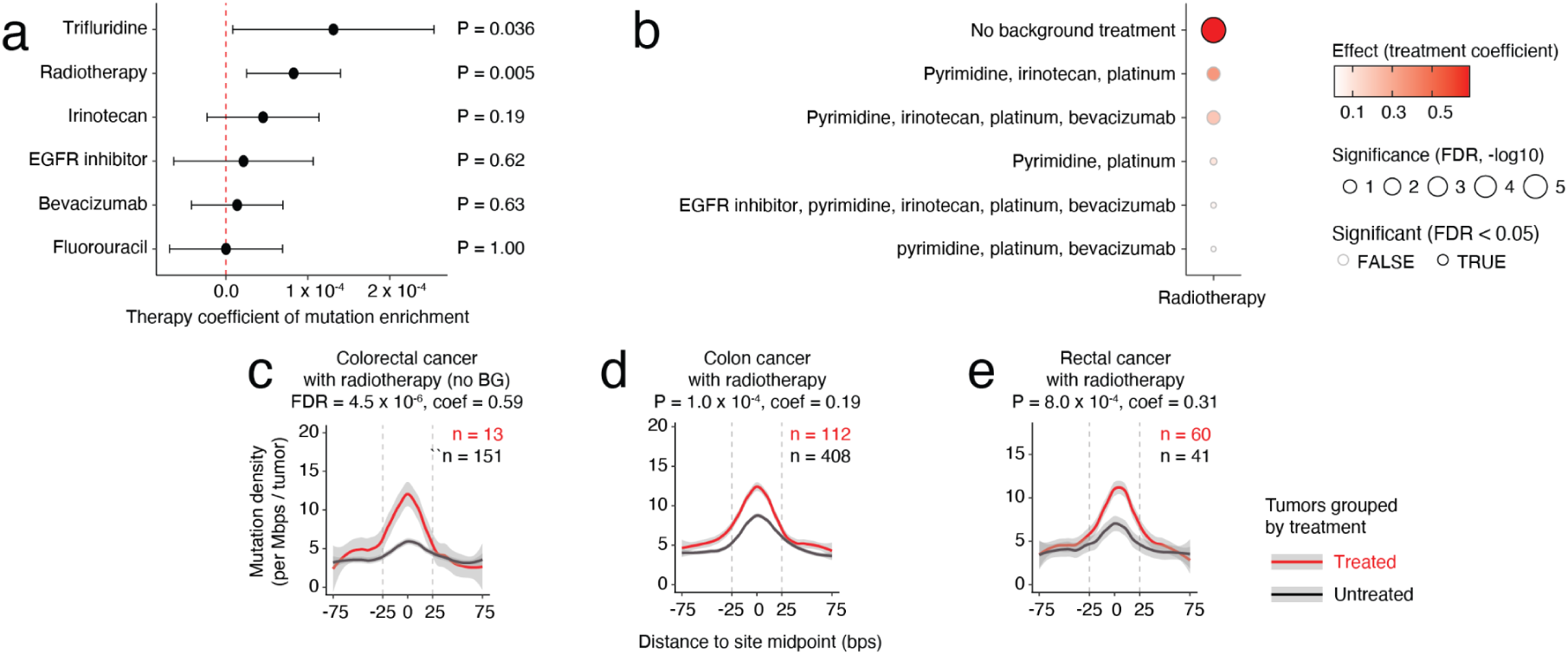
Radiotherapy and trifluridine treatments are associated with increased mutation enrichment at CTCF binding sites in metastatic colorectal cancer. **(a)** Independent effects of major colorectal cancer therapies on CBS mutation enrichment. A multivariable linear model was used to associate CBS mutation enrichment with overlapping treatment exposures. The point-range plot shows regression coefficients with 95% confidence intervals; positive coefficients indicate increased enrichment. **(b)** Radiotherapy-associated CBS mutation enrichment across background treatment contexts. *P*-values corresponding to treatment coefficients were computed using RM2 and adjusted for multiple testing using FDR. Pyrimidine: Pyrimidine analog treatments. Platinum: Platinum-based chemotherapy. **(c)** Stringent comparison of tumors with no background (BG) treatment. Treated tumors were exposed to only radiotherapy treatment while the treatment-naive (untreated) tumors were not exposed to any treatment. **(d-e)** Radiotherapy-associated CBS mutation enrichment analyzed separately in colon (d) and rectal (e) cancers, confirming the association in both subcohorts. In panels (c-e), the y-axis shows aggregated mutation density across all CBS. Lines show LOESS-smoothed mutation density (span = 0.4) and the shading indicates standard error. Sample sizes and significance of CBS mutation enrichment relative to flanks are shown for treated (red text) and untreated groups (black text). The x-axis denotes sequence distance (bp) from CBS midpoints. Gray dotted lines indicate CBS boundaries (50 bp).

To further assess whether the association between radiotherapy and CBS mutation enrichment was robust to treatment context, we stratified radiotherapy-treated colorectal tumors by the combinations of other therapies the patients received (**Figure 2b, Table S5**). Notably, tumors that received radiotherapy as their only documented treatment exhibited significantly increased CBS mutation enrichment compared to tumors with no recorded treatment exposure (FDR = 4.5 x 10^-6^ from RM2; **Figure 2c**). This comparison provides evidence that radiotherapy exposure itself is associated with localized mutation enrichment at CBS, independent of other therapies. Across other background treatment groups, radiotherapy-treated tumors generally showed higher CBS mutation enrichment than tumors without radiotherapy exposure (**Figure 2b**). However, most individual comparisons were limited by sample sizes within each subgroup and did not reach statistical significance after multiple-testing correction.

Radiotherapy is more commonly administered to rectal than colon cancers [30], despite these tumor types sharing highly similar genomic and transcriptomic features [31,32], raising the possibility that tumor location could confound the observed association. We analyzed colon and rectal cancers separately and observed significant radiotherapy-associated CBS mutation enrichment in both the colon cancer cohort (*P* = 1.0 x 10⁻⁴) and the rectum cancer cohort (*P* = 8.0 x 10⁻⁴) (**Figure 2d-e**). To further minimize potential confounding effects, we performed a stringent comparison between colorectal tumors exposed only to radiotherapy (n = 13) and treatment-naive tumors matched for sample size and rectal cancer proportion. Even under this conservative design, tumors treated with radiotherapy alone showed significantly higher CBS mutation enrichment than fully untreated tumors (*P* = 0.032 from RM2; **Figure S2**), indicating that the association is not driven by tumor location.

### Chromatin context and DNA-binding motifs shape mutation enrichment at CTCF binding sites

Focusing on radiotherapy-treated colorectal cancers, we next examined whether local genomic and chromatin features modulate mutation enrichment at binding sites. We annotated constitutively active CBS with multiple properties, including the presence of a canonical CTCF DNA-binding motif, replication timing, expression level of nearby genes, chromatin loop anchoring, and topologically associated domain (TAD) context. CBS were stratified by each feature, and RM2 was applied within strata to assess radiotherapy-associated changes in mutation enrichment relative to flanking regions. Across these analyses, mutation enrichment and its association with radiotherapy were strongly influenced by DNA motif presence, replication timing, and transcriptional activity, whereas chromatin loop anchoring and TAD context showed little or no effect (**Figure 3a, Table S6**).

**Figure 3.**
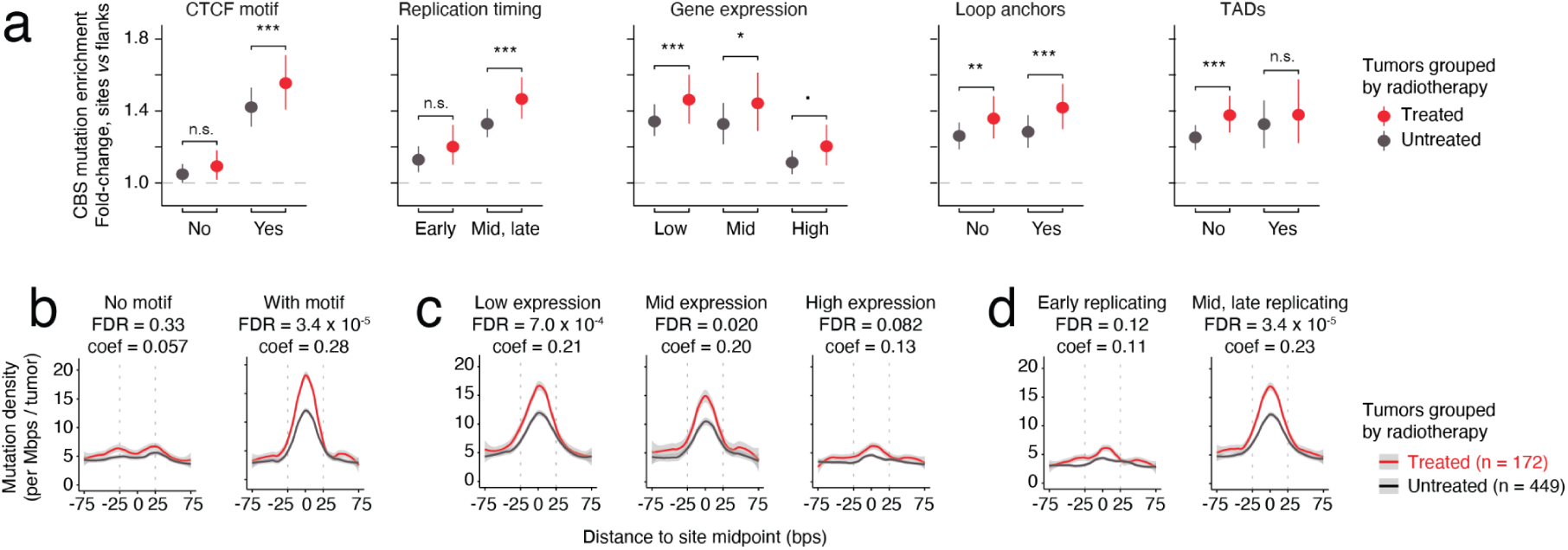
DNA motif presence and local chromatin context modulate mutation enrichment at binding sites. **(a)** CBS mutation enrichment in radiotherapy-treated and untreated colorectal tumors stratified by genomic context of CBS. Pointrange plots show fold-change relative to flanking regions (95% CI). FDR-corrected significance of therapy coefficients from RM2 are shown. Asterisks show P-values (*** P < 0.001; ** P < 0.01; * P < 0.05; ‧ P < 0.1). (**b-d)** CBS mutation enrichment profiles stratified by (b) motif presence, (c) nearby gene expression, and (d) replication timing. In panels (b-d), the y-axis shows aggregated mutation density across all CBS. Lines show LOESS-smoothed mutation density (span = 0.4) and the shading indicates standard error. Sample sizes are shown for treated (red text) and untreated groups (black text). The x-axis denotes sequence distance (bp) from midpoints of CBS. Gray dotted lines indicate boundaries of CBS used in this analysis (50 bp).

The presence of a canonical CTCF DNA-binding motif emerged as the strongest determinant of mutation enrichment at CBS (**Figure 3a-b**). CBS containing the canonical motif showed clear mutation enrichment relative to flanking regions in both radiotherapy-treated and untreated tumors. Importantly, radiotherapy exposure was associated with a further increase in mutation enrichment at motif-containing CBS (*P* = 3.4 x 10^-5^ from RM2). In contrast, CBS lacking the canonical motif showed little or no mutation enrichment in either treatment group and no detectable radiotherapy-associated effect. These results indicate that mutation enrichment is highly concentrated at CBS with CTCF DNA-binding motifs, both at baseline and following radiotherapy exposure, extending prior observations of CBS-focused mutagenesis [19]. Because motif presence is a proxy for direct and stable CTCF-DNA interaction [33], this pattern is consistent with the notion that sustained protein binding may constrain local DNA repair and thereby increase the likelihood that radiation-induced damage is fixed as somatic mutations.

Transcriptional activity and replication timing further modulated mutation enrichment at CBS and its association with radiotherapy exposure. To study transcriptional associations, we grouped CBS based on low, middle or high gene expression. In both low and mid-expression groups, radiotherapy exposure was associated with a significant increase in CBS mutation enrichment (lowly-expressed CBS, *P* = 7.5 x 10^-4^; mid-expressed CBS, *P* = 0.02; RM2) (**Figure 3a, c**). Similarly, CBS in late-replicating genomic regions showed a significantly elevated mutation enrichment in the genomes of tumors having prior radiotherapy exposure (*P* = 3.4 x 10^-5^; **Figure 3a, d**). In both comparisons, both treated and untreated tumor subsets showed mutational enrichments in CBS compared to flanking genomic regions (**Figure 3a, c-d**). In contrast, CBS near highly expressed genes or in early-replicating genomic regions showed little or no radiotherapy-associated effect and no mutation enrichment in CBS overall. Because DNA repair efficiency is reduced in late-replicating regions and transcription-coupled repair operates primarily in actively transcribed regions [34,35], these patterns are consistent with radiotherapy-associated mutagenesis preferentially affecting CBS located in genomic contexts with limited repair activity. Overall, these results indicate that CBS mutation enrichment, and its enhancement by radiotherapy, is strongly shaped by DNA motif presence and local chromatin context.

### Treatment-associated mutation enrichment at CTCF binding sites is dominated by SBS5

We next examined the mutational spectrum underlying treatment-associated CBS mutation enrichment. In metastatic colorectal cancer, the increased mutation burden at CBS was overwhelmingly driven by single nucleotide variants (SNVs), which accounted for the large majority of mutations at CBS (94%). Indeed, SNVs were highly enriched in sites relative to flanking sequences, with radiotherapy exposure associated with a further increase in mutation enrichment (FDR = 3.9 x 10^-5^ from RM2; **Figure 4a, Table S7**). In contrast, indels in CBS were depleted relative to flanking regions (*P* < 0.0001) and no association with radiotherapy exposure with indel mutations was identified at CBS (**Figure 4b**), despite prior reports linking ionizing radiation to small deletions [27]. We therefore focused subsequent analyses on SNVs.

**Figure 4.**
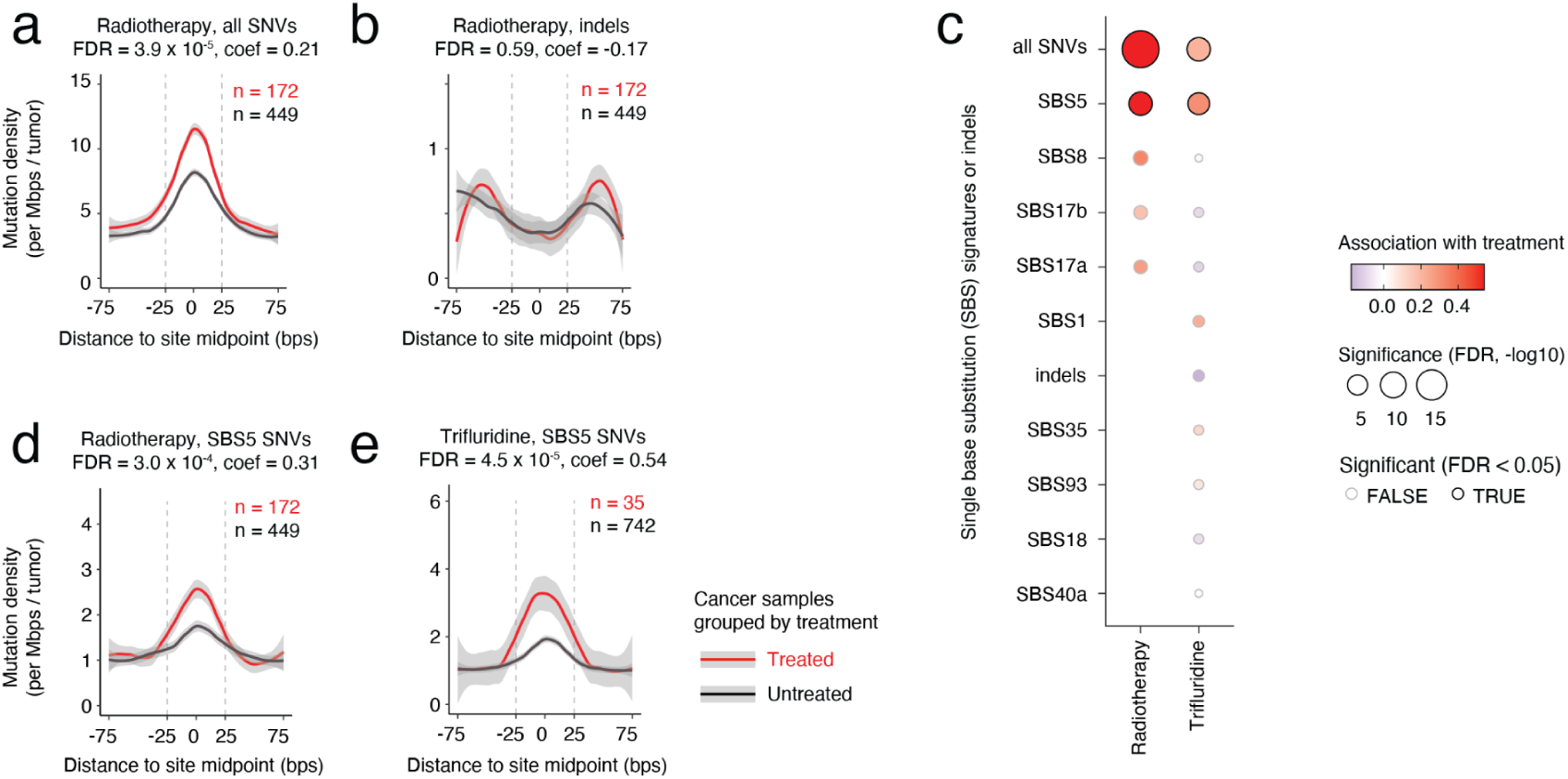
Treatment-associated mutation enrichment at CTCF binding sites is predominantly attributed to SBS5. (a-b) Contribution of SNVs (a) and indels (b) to CBS mutation enrichment in radiotherapy-treated and untreated colorectal tumors. P-values from RM2 reflect therapy contributions to CBS mutation enrichment. **(c)** Analysis of SBS signatures at CBS in the context of treatment exposures in colorectal cancer. Each dot represents an SBS signature tested for treatment-associated changes in CBS mutation enrichment. Dot color indicates the direction and magnitude of treatment effects on CBS enrichment, where positive values indicate increased CBS mutation enrichment in treated tumors. Dot size reflects FDR-adjusted significance from RM2. **(d-e)** In both panels, the Y-axis shows aggregated mutation density across all CBS (*i.e.*, mutations per Mbps per sample). Lines show LOESS-smoothed mutation density (span = 0.4) and the shading indicates standard error. The x-axis denotes sequence distance (bp) from midpoints of CBS. Gray dotted lines indicate boundaries of CBS used in this analysis (50 bp).

To determine which mutational processes underlie treatment-associated mutation enrichment at CBS, we stratified SNVs by COSMIC mutational signatures of single base substitutions [11] (SBS) (**Figure 4c**). Across signatures commonly observed in colorectal cancer, SBS5 showed a pronounced increase in CBS mutation enrichment in association with radiotherapy or trifluridine exposure (FDR = 3.0 x 10^-4^ and FDR = 4.5 x 10^-5^, respectively from RM2; **Figure 4c-e**). SBS5 is a broadly distributed signature, lacking strong trinucleotide preferences, that has been described as a clock-like signature in multiple cancer types [11,36]. Although its aetiology remains incompletely understood, SBS5 has been linked to the accumulation of endogenous DNA damage and imperfect DNA repair [37]. The preferential association of treatment-related CBS mutation enrichment with SBS5 is therefore consistent with therapy exposure contributing to increased mutation accumulation at sites with constrained DNA repair. In contrast, other mutational signatures prevalent in colorectal cancer, including SBS8, SBS17a, and SBS17b, showed mutation enrichment at CBS in both treated and untreated tumors but did not exhibit any association with radiotherapy or trifluridine exposure (**Figure S3**). Thus, while multiple endogenous mutational processes contribute to baseline CBS mutation burden in colorectal cancer, the treatment-associated increase in CBS mutation enrichment is primarily captured by SBS5.

Because SBS5 activity has been reported to correlate with age in several cancer types [36], we next assessed whether age differences between treatment groups could confound the observed association between SBS5-associated CBS mutations and therapy exposure. Patient age at biopsy was comparable between radiotherapy-treated and untreated colorectal tumors (**Figure S4a**), as well as between trifluridine-treated and untreated tumors (**Figure S4b**). This is consistent with prior reports showing that SBS5 is not strongly correlated with age in primary colorectal cancer [36]. Together, these observations indicate that age is unlikely to account for the increased SBS5-associated mutation enrichment at CBS observed in radiotherapy- and trifluridine-exposed tumors, supporting a treatment-associated origin of this effect.

### DNA damage response gene alterations modify radiotherapy-associated mutation enrichment at CTCF binding sites

Alterations in genes involved in genome maintenance and DNA damage response are common in colorectal cancer and can influence how tumors respond to genotoxic stress. We therefore asked whether such alterations modify radiotherapy-associated mutation enrichment at CBS. We stratified radiotherapy-treated colorectal tumors by the mutation status of recurrently-altered driver genes annotated in the HMF dataset [28]. For each driver gene, we then compared radiotherapy-associated CBS mutation enrichment between driver-mutant and wildtype tumors using RM2.

We detected six driver genes whose alterations associated with differences in CBS mutation enrichment within the subset of radiotherapy-treated colorectal tumors (FDR < 0.05; **Figure 5a, Table S8**). Most prominently, *BRCA2* alterations, observed in ten tumors, were associated with a substantially higher degree of CBS mutation enrichment. These *BRCA2*-mutant tumors exhibited greater CBS mutation enrichment than 162 *BRCA2*-wildtype tumors following radiotherapy exposure (1.62-fold *vs.* 1.33-fold enrichment; FDR = 4.4 x 10⁻⁵; RM2; **Figure 5b**). BRCA2 plays a central role in homologous recombination mediated DNA repair, and BRCA2 deficiency compromises DNA double-strand break repair [38,39]. The increased CBS mutation enrichment observed in *BRCA2*-mutant, radiotherapy-treated tumors is therefore consistent with impaired repair of radiation-induced damage, increasing the likelihood that unresolved DNA lesions are fixed as mutations at sites where DNA repair is already constrained.

**Figure 5.**
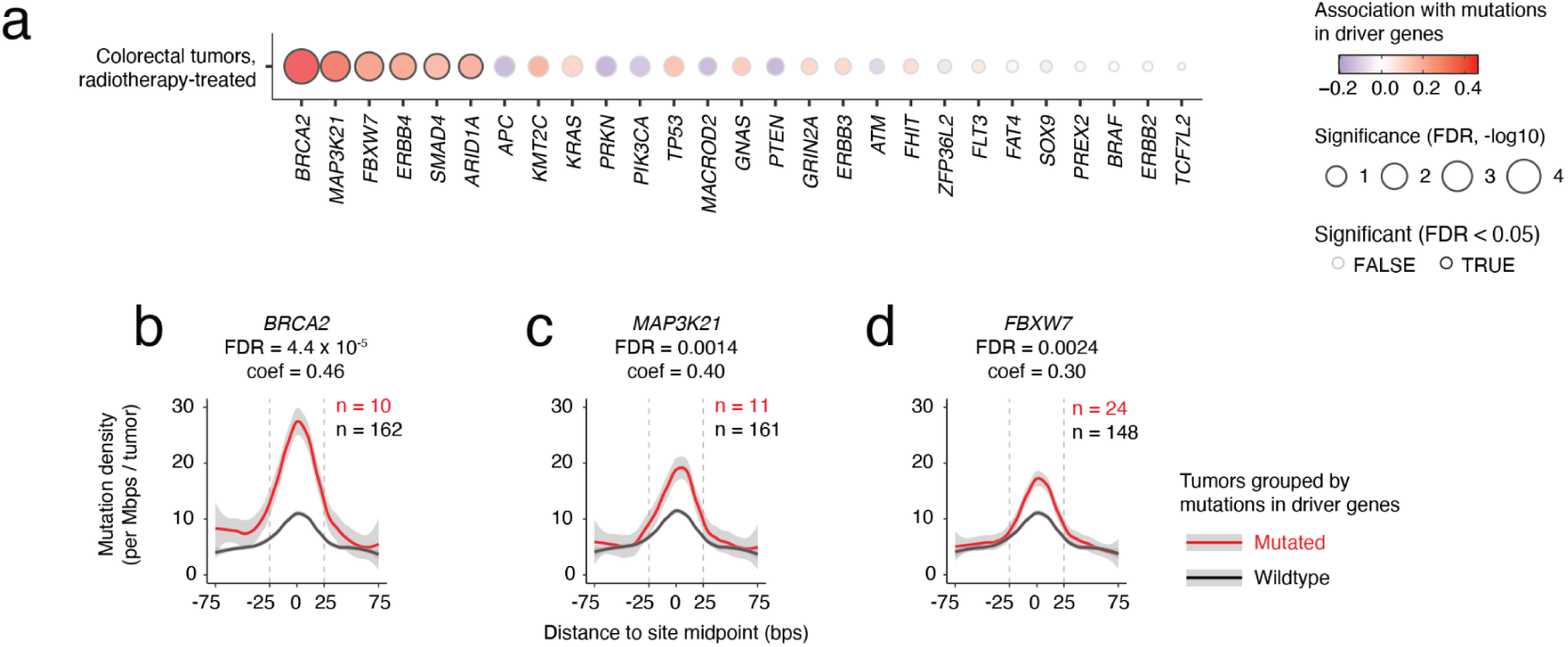
Alterations in DNA damage response genes modify radiotherapy-associated mutation enrichment at CTCF binding sites. **(a)** Association between driver mutation status and CBS mutation enrichment in radiotherapy-treated colorectal tumors. Each dot in the plot represents a driver gene tested, measuring the contribution of driver mutation status to CBS mutation enrichment, where positive values indicate increased CBS mutation enrichment in driver-mutant tumors and dot size reflects significance. Statistical significance was computed using RM2 and P-values were adjusted for multiple testing using FDR. (**b-d)** CBS mutation enrichment profiles in radiotherapy-treated colorectal tumors stratified by driver mutation status of top genes (*BRCA2*, *MAP3K21*, *FBXW7*). In all three panels, the Y-axis shows aggregated mutation density across all CBS (*i.e*., mutations per Mbps per sample). Lines show LOESS-smoothed mutation density (span = 0.4) and the shading indicates standard error. The x-axis denotes sequence distance (bp) from midpoints of CBS. Gray dotted lines indicate boundaries of CBS used in this analysis (50 bp).

In addition to *BRCA2*, alterations in several other driver genes were associated with increased CBS mutation enrichment in the genomes of radiotherapy-treated colorectal tumors (**Figure 5a**). Notably, mutations in *MAP3K21* and *FBXW7* were associated with higher CBS mutation enrichment compared to wildtype tumors (**Figure 5c-d**). Both genes have been implicated in DNA damage sensing and the regulation of non-homologous end joining [40–42], a major pathway for the repair of double-strand breaks induced by ionizing radiation [43]. Three additional genes (*ERBB4*, *ARID1A*, *SMAD4*) also showed significant associations in the systematic analysis, although the differences in CBS mutation enrichment between wildtype and mutant tumors were modest (**Figure S5**). Overall, these results indicate that radiotherapy-associated mutation enrichment at CTCF binding sites is shaped not only by local chromatin context but also by tumor-specific defects in DNA damage response pathways, pointing to a convergence between therapy-induced DNA damage and impaired repair at key regulatory elements.

## Discussion

In this study, we systematically evaluated whether genotoxic cancer therapies are linked to localized mutation enrichment at CTCF binding sites (CBS) in metastatic tumors. In metastatic colorectal cancer, radiotherapy and trifluridine exposure were associated with increased CBS mutation enrichment, and these associations persisted after accounting for overlapping treatment histories. These findings indicate that cancer therapies can reshape mutagenesis at architectural regulatory elements and suggest that the genomic consequences of treatment are influenced by chromatin context and DNA repair capacity. Although the strongest associations were observed in colorectal cancer, CBS mutation enrichment is also present in tumors not exposed to radiotherapy, and therapy exposure appears to amplify this baseline pattern, suggesting a contribution that is complementary to intrinsic differences between cancer types.

Therapy-associated CBS mutation enrichment was further associated with functional characteristics of non-coding DNA. CBS containing a canonical CTCF motif showed the strongest mutation enrichment, consistent with stable CTCF-DNA occupancy and the influence of epigenetic context on binding [33]. Because transcription factor binding has been shown to impair nucleotide excision repair [14], sustained CTCF occupancy may constrain local repair accessibility, increasing the likelihood that DNA damage is fixed as mutation at these sites. Consistent with the idea that CTCF sites are hotspots of DNA break-associated mutagenesis, we previously observed that CBS co-bound by the topoisomerase TOP2B exhibit marked enrichment of SNVs in cancer genomes, implicating double-strand break dynamics and repair processes at these regulatory elements [21]. These findings support a model in which therapy-associated mutagenesis preferentially occurs at regulatory elements where protein occupancy limits repair efficiency.

Replication timing and transcriptional activity further contextualize these findings. Late-replicating regions are known to exhibit higher baseline mutation rates, partially due to reduced DNA repair efficiency and replication-associated stress [44,45]. Regional variation in DNA repair pathways, including DNA mismatch repair, also contributes to variation in genome-wide mutation process activity [35]. Transcription-coupled and global nucleotide excision repair pathways play central roles in the removal of helix-distorting lesions and the prevention of mutagenesis across the genome [46]. In our data, radiotherapy-associated CBS mutation enrichment was most evident within late-replicating regions and near lowly expressed genes, consistent with reduced repair activity in these genomic contexts. Together, these observations support a model in which therapy-induced DNA damage is more likely to be fixed as mutation at repair-constrained CTCF binding sites.

Consistent with this repair-constrained model, alterations in multiple DNA damage response genes further amplified radiotherapy-associated CBS mutation enrichment. *BRCA2*-mutant tumors exhibited higher CBS mutation enrichment following radiotherapy, in line with the established role of BRCA2 in homologous recombination-mediated repair of DNA double-strand breaks [38,39]. Similarly, mutations in *MAP3K21* and *FBXW7*, which are implicated in DNA damage signaling and regulation of non-homologous end joining [40–42], were associated with increased CBS mutation enrichment in radiotherapy-treated tumors. Together, these findings are consistent with the idea that compromised double-strand break repair capacity increases the likelihood that radiation-induced lesions are fixed as mutations at CTCF binding sites, where repair accessibility is already limited.

Mutational signature analysis indicated that treatment-associated CBS mutation enrichment was primarily captured by SBS5. Although SBS5 lacks strong trinucleotide specificity and is broadly observed across tumor types [11,47], it has been linked to endogenous DNA damage and imperfect DNA repair [36,37]. The association of radiotherapy- and trifluridine-related CBS enrichment with SBS5 is therefore consistent with increased fixation of DNA lesions at repair-constrained sites rather than the emergence of a distinct, therapy-specific mutational signature. While radiation-associated mutational patterns have been described in experimental systems and specific tumor contexts [3,27,48–51], a consistent and universally accepted radiation signature in human cancers remains elusive. The relatively featureless profile of SBS5 is compatible with the stochastic fixation of DNA lesions arising from ionizing radiation-induced damage, including single-strand breaks that may be converted into SNVs when improperly repaired [52].

CBS serve as anchors of chromatin loops and topologically associated domains, and disruption of CTCF-mediated insulation has been shown to alter promoter-enhancer interactions and activate proto-oncogenes through changes in chromosomal neighborhood structure [20,23]. Somatic mutations at CTCF insulators have also been identified as candidate non-coding driver alterations with regulatory impact in cancer genomes [53]. In addition, we previously demonstrated that CBS co-bound by the topoisomerase TOP2B and architectural factors such as CTCF and RAD21 exhibit marked enrichment of small mutations, especially at known cancer driver loci [21]. Because mutation enrichment in the present study was the strongest at motif-containing CBS, where stable CTCF occupancy is expected, therapy-associated mutations may perturb CTCF binding and subtly reshape local chromatin architecture. Such alterations could influence nearby gene expression or regulatory domain boundaries, contributing to increased molecular heterogeneity within treated tumors.

This study has several limitations. Treatment exposures prior to and following development of metastatic disease are complex and frequently correlated because patients commonly receive multi-agent regimens and successive lines of therapy. Although joint modeling was used to estimate independent effects, residual confounding cannot be excluded. In addition, because this cohort consists of metastatic tumors obtained after therapy, it represents a selected population of cancers that persisted or recurred following treatment. The observed mutation enrichment therefore reflects evolutionary processes in treatment-surviving disease and may not fully capture mutational consequences in tumors that respond durably to therapy. The dataset also lacks matched pre-treatment and post-treatment samples and treatment dosage information, precluding direct causal attribution of CBS mutations to specific therapies. Finally, statistical power varied across pairs of cancer types and therapies, and was reduced in subgroup analyses of radiotherapy-treated tumors stratified by driver mutation status, which may limit detection of smaller effect sizes.

Radiotherapy remains a central component of treatment across many cancer types for curative or palliative intent, and trifluridine/tipiracil represents a more recently introduced therapy for refractory metastatic colorectal cancer that improves survival in previously treated patients [54]. Our findings indicate that, beyond their intended cytotoxic effects, genotoxic therapies leave measurable mutational imprints at architectural regulatory elements of the non-coding genome. This study extends the genomic consequences of cancer treatment to higher-order genome organization by linking therapy exposure to localized CBS mutagenesis and demonstrating its modulation by chromatin context and DNA repair capacity. A deeper understanding of therapy-associated mutagenesis in regulatory regions may refine models of tumor evolution in treated cancers and inform future studies of genomic stability in this setting.

## Methods

### Whole-genome sequencing (WGS) data

WGS data for metastatic tumors in hg19 were obtained from the Hartwig Medical Foundation (HMF) cohort [28] (downloaded on 2025-07-10). All sequencing and primary processing were performed by HMF under locally approved Institutional Review Board protocols with written informed consent, as described previously [28]. This secondary analysis was approved by the University of Toronto Research Ethics Board (protocol 37521). We used HMF-provided somatic variant calls, driver annotations, RNA-seq profiles (when available), and clinical metadata including pre-biopsy treatment history, age at biopsy, and metastatic biopsy site [28]. Tumors were grouped by primary cancer type and cancer types with fewer than 25 samples or of unknown primary type were excluded. When multiple biopsies per patient were available, we selected one sample based on treatment information availability. To reduce the influence of extreme hypermutators, tumors with >90,000 somatic mutations were excluded. Colon and rectal adenocarcinomas were analyzed together as colorectal tumors for primary analyses and we also confirmed findings in colon and rectal tumors separately.

### Treatment information and clinical annotations

Pre-biopsy treatment histories were curated to harmonize treatment names, aliases, and regimen components. Because related therapies such as those within the same drug class may induce similar mutational patterns [3], we analyzed both individual treatments and predefined treatment groups (**Table S2**). Treatments were grouped using the hierarchy from the NCI Thesaurus (v25.10d) [55]. For grouped analyses, a tumor was labeled “treated” if the patient received any therapy within the group prior to biopsy, and “untreated” otherwise. Patient age at biopsy was estimated from birth year and biopsy year. Age could not be derived for 881 patients due to missing birth year and/or biopsy year; these samples were excluded from analyses requiring age.

### CTCF binding sites (CBS)

CBS were defined as the 32,442 constitutively active binding sites from our prior work [19]. Briefly, previously preprocessed CTCF ChIP-seq peaks from 70 ENCODE cell lines spanning diverse tissues were aggregated [29,56]. The peaks were divided into 5 bins based on the number of cell lines with active CTCF binding. Bin 1 was bound by CTCF in 52-70 cell lines (median 67) and was defined as constitutively active sites. This subset of sites showed the strongest and most consistent mutation enrichment in primary tumors in our prior analysis [19]. As a quality check, we confirmed that mutation enrichment in the HMF metastatic cohort was likewise concentrated in this bin (**Figure S1**).

### Evaluating local mutation enrichment in CBS

We used RM2 (v1.0.5), a negative binomial regression framework for testing localized mutation enrichment variation in a class of genomic elements relative to local background [19]. For each site, we defined the element as the central sequence of 50 bp (±25 bp from peak centre) and used the immediately flanking regions of matched size (50 bp on either side) as local background. RM2 models mutation counts with trinucleotide context and megabase-scale background mutation burden as covariates. A CBS-specific enrichment effect is supported when the model including a site indicator fits significantly better than the model without it, based on a likelihood ratio test. To compare groups of tumors based on genomic or clinical annotation (e.g., treatment exposure, driver mutation status), we included an interaction term between group label and the site indicator (*i.e.*, CBS *vs.* flanking genomic sequences) as defined in RM2. Significance was assessed by likelihood-ratio (chi-square) testing of models with vs without the interaction. A significant interaction indicates that the group modifies CBS mutation enrichment relative to local background (*i.e.*, alters the CBS-to-flank mutation enrichment ratio). P-values from RM2 were adjusted for multiple testing correction using Benjamini-Hochberg false discovery rate (FDR) [57] and significant results were selected (FDR < 0.05).

### Analysing treatment combinations associated with CBS mutation enrichment in colorectal cancer

To estimate the independent contributions of overlapping therapies in colorectal cancer, we fitted a multivariable linear regression model with binary treatment exposures as predictors and the fraction of somatic mutations occurring within CBS for each tumor, defined as the number of CBS mutations (SNVs, indels) divided by total number of somatic mutations in that tumor. Six treatment exposures were included (trifluridine, radiotherapy, irinotecan, bevacizumab, fluorouracil, anti-EGFR). Coefficients and 95% confidence intervals were estimated from the regression models and two-sided t-tests were used to test whether each coefficient differed from zero.

### Single base substitution (SBS) signatures and CBS mutations

The SigProfilerAssignment method and R package (v0.0.23) [58] was used to estimate the contribution of SBS signatures [11] in each whole cancer genome using SBS signature definitions from the COSMIC database (v3.4) [59]. For each SNV trinucleotide category in a sample, SigProfilerAssignmentR provides the posterior probability of an SNV being generated by each signature given its trinucleotide context. Mutations were then assigned to the SBS signature with the highest probability given their trinucleotide, similarly to earlier studies [60–62]. To analyze specific mutational signatures, RM2 was used on individual subsets of SNVs assigned to each of the SBS signatures, excluding infrequent signatures having fewer than 100 SNVs in CBS and their flanking regions.

### Stratifying CBS by genomic and functional characteristics

To functionally characterize mutation enrichment at CBS with respect to radiotherapy treatment, CBS were annotated by canonical DNA motif presence, replication timing, nearby gene expression, overlap with chromatin loop anchors, and overlap with TAD boundaries. Presence of canonical DNA-binding motifs of CTCF at CBS were obtained from the study by Lee *et al*. [17], which were determined using the FIMO method [63] with the motif MA0139.1 from the JASPAR database [64], selecting sites with significant matches (FDR < 0.01). Replication timing profiles of the colorectal cancer cell line HCT116 were obtained from a previous study [65] (GEO: GSE158008). Genomic bins of 1 kbp were grouped as early-replicating (top 25%), middle-replicating (middle 50%) and late-replicating regions (bottom 25%) based on weighted average (WA) scores. Subsets of CBS that either overlapped several types of regions (29 sites) or lacked WA (1101 sites) were excluded. Due to fewer CBS in late-replicating regions (**Figure S6**), CBS in mid-replicating and late-replicating regions were grouped and compared to CBS in early-replicating regions in relevant analyses of CBS-based mutational processes. For transcriptional activity at CBS, we used matching RNA-seq data of 391 colorectal tumors from HMF [28], which included protein-coding genes and lncRNAs. We computed median expression values for each gene across the samples and then divided the genes into three equal groups as high, middle, or low expression. CBS were next assigned to genes based on genomic windows of 3 kbp. CBS were annotated as high expression if they were adjacent to at least one highly-expressed gene. Sites that are adjacent to at least one mid-expression gene but not any high-expression genes were annotated as mid-expressed, and otherwise annotated as lowly-expressed. Next, chromatin loop anchors from two colorectal cancer cell lines (HT29, LoVo) were retrieved from a previous promoter-capture HiC study [66]. TAD boundary information from human embryonic stem cells (H1) and lung fibroblast cell lines (IMR90) was retrieved from a previous study [67], assuming that TAD boundaries are largely conserved between cell types [67] and thus appropriate for analysis of mutations in colorectal cancer. CBS were annotated at loop anchors and TAD boundaries based on coordinate overlaps with CBS based on constitutive CTCF ChIP-seq peaks derived from ENCODE [29]. For each subset of feature-stratified CBS, RM2 was used to analyze CBS-to-flank mutation enrichment in samples grouped by radiotherapy status. P-values from RM2 were adjusted for multiple testing correction using Benjamini-Hochberg FDR and significant results were selected (FDR < 0.05).

### Analysing driver mutations associated with CBS mutation enrichment in colorectal cancer

Driver mutation status of each sample was obtained from HMF [28], which considers protein-coding SNVs and indels, splicing disruptions, mutation hotspots, biallelic inactivation, dNdS, and copy number alterations, as described previously [68]. Driver gene modifier analyses were restricted to radiotherapy-treated colorectal tumors while trifluridine treatment was not separately analysed due to limited sample size. We designated samples as mutated in a driver gene if they had any functional mutations in the gene based on HMF annotations, and wildtype otherwise. We analyzed driver genes with at least 10 samples in both mutated and wildtype groups. RM2 was used to compare CBS mutation enrichment in samples grouped by driver status (mutated *vs* wildtype). P-values from RM2 were adjusted for multiple testing correction using Benjamini-Hochberg FDR and significant results were selected (FDR < 0.05).

## Data availability

Processed data are available as supplementary datasets. Input datasets from Hartwig Medical Foundation (HMF) representing WGS, transcriptomics, and metadata annotations of cancer samples of multiple cancer types are controlled-access and can be provided by HMF pending scientific review and a completed material transfer agreement. Requests for these datasets should be submitted to the Hartwig Medical Foundation. Intermediate data files cannot be shared due to the use of controlled-access datasets from HMF.

## Code availability

Publicly available software packages and common statistical methods were used to process and analyze the data as described in the Methods section.

## Acknowledgments

We would like to thank Diogo Pellegrina, Nina Adler and Mykhaylo Slobodyanyuk for valuable discussions. This work was partially supported by the Canadian Institutes of Health Research (CIHR) Project Grants (PJT-162410, PJT-197925) and Catalyst Grant (DV1-197665) to J.R., the Investigator Award to J.R. from the Ontario Institute for Cancer Research (OICR), and the New Investigator Award of the Terry Fox Research Institute (TFRI-PROJECT-1095) to J.R. A.B. was supported by the Ontario Graduate Scholarship (OGS). K.C. was partially supported by fellowships from the Medical Biophysics Department at University of Toronto. Research in the B.H. Lok laboratory is funded by the Terry Fox Research Institute, Canada Foundation for Innovation, Canadian Institutes of Health Research, National Institute of Health/National Cancer Institute (U01CA253383), Clinical and Translational Science Center at Weill Cornell Medical Center, MSKCC (UL1TR00457). Funding to OICR is provided by the Government of Ontario. This publication and the underlying study have been made possible partly based on the data that Hartwig Medical Foundation has made available to the study.

## Author Contributions

K.C. performed formal analysis and data visualisation with input from J.R.. K.C. and J.R. interpreted the data. J.M. contributed to data visualisation. K.C. and Z.K. pre-processed clinical data. Z.K. pre-processed RNA-seq data. K.C., J.M., Z.K., A.B., B.L., T.P. and J.R. contributed to methodology and data interpretation. J.R. conceptualized the idea, supervised the project, and acquired funding. K.C. and J.R. wrote the manuscript. All coauthors reviewed, edited and approved the final manuscript.

## Supplementary tables

**Table S1. RM2 results of treatment-associated mutation enrichment.** This table shows the results of the initial systematic analysis of cancer types and therapy types as presented in Figure 1B.

**Table S2. Coordinates and properties of CTCF binding sites**. This table shows genomic locations of constitutively active CTCF binding sites based on the human genome version hg19. Functional properties including presence of CTCF binding motif, DNA replication timing, gene expression association, presence of loop anchors and TAD involvement are also listed.

**Table S3. Cancer treatments grouped by similarity.** This table shows the major cancer therapies applied in the HMF cohort. Therapies are arranged into groups of similar therapies according to the NCI Thesaurus.

**Table S4. Overview of the colorectal cancer cohort**. This table shows the summary statistics of colorectal cancer patients in the HMF cohort that were included in the analyses of radiotherapy associations and mutation enrichment at CBS.

**Table S5. RM2 results of radiotherapy-associated CBS mutation enrichment in background treatment matched samples.** This table shows radiotherapy-associated mutation enrichment increase in the colorectal cancer cohort. Samples were grouped by their co-treatment history (i.e., treatments received other than radiotherapy). The data are presented in Figure 2B.

**Table S6. RM2 results of radiotherapy-associated mutation enrichment in CBS grouped by genomic context**. This table shows radiotherapy-associated mutation enrichment increase in the colorectal cancer cohort across subsets of CTCF binding sites, grouped by their genomic features including CTCF binding motif, DNA replication timing, gene expression association, loop anchor involvement and TAD involvement as presented in Figure 3A.

**Table S7. RM2 results of treatment-associated enrichment of COSMIC mutational signatures in CBS**. This table shows mutational signatures associated with radiotherapy and trifluridine in the colorectal cancer cohort as presented in Figure 4C.

**Table S8. RM2 results of driver gene alteration-associated CBS mutation enrichment in radiotherapy-treated colorectal cancer**. This table shows RM2 results quantifying the association between driver gene alteration status and CBS mutation enrichment within radiotherapy-treated colorectal tumors, comparing driver-mutant against wildtype tumor subgroups for each tested gene, as presented in Figure 5A.

