## Supplementary Figures for "Therapy-associated mutagenesis at CTCF binding sites is shaped by chromatin context and DNA repair capacity"

### Supplementary information

Kevin CL Cheng *et al.* (2026)

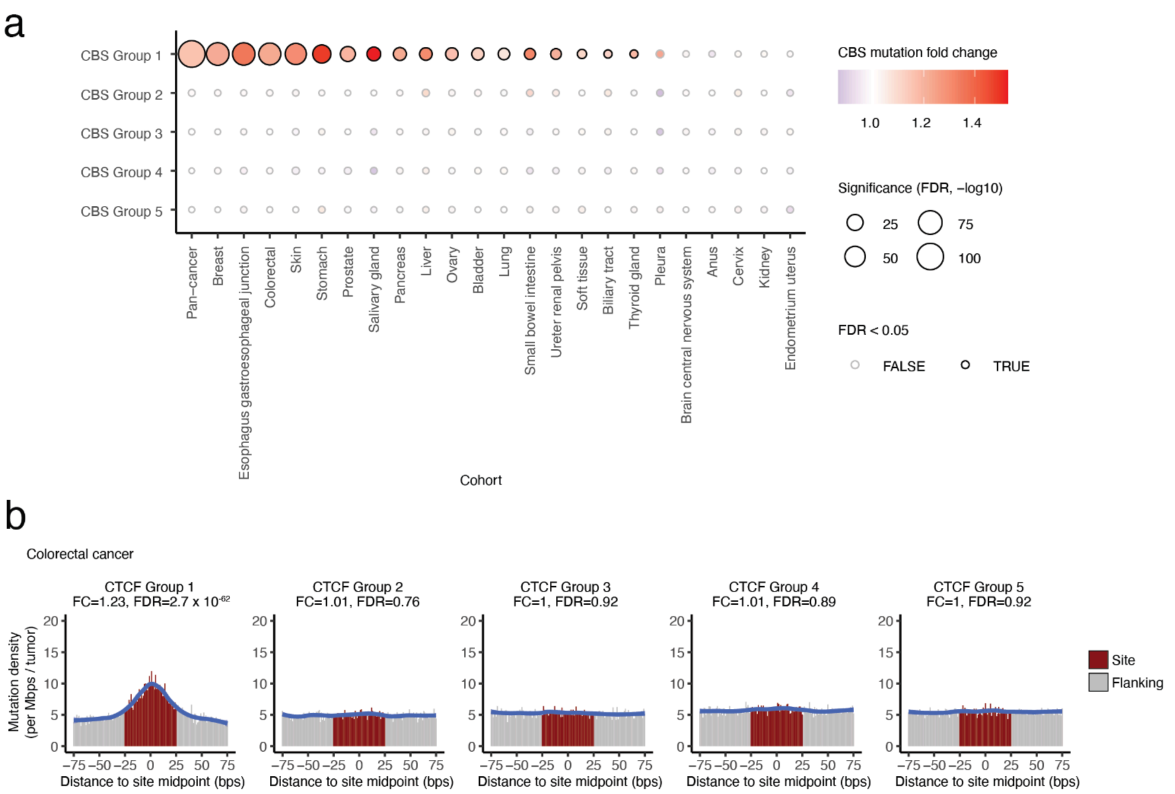


Figure S1. Mutation enrichment of CTCF binding sites (CBS) in metastatic cancer grouped by cross-tissue CTCF binding activity.

CBS were grouped by binding activity across 70 ENCODE cell types. Group 1 CBS were constitutively bound by CTCF across most tissue types whereas Groups 2-5 show progressively more tissue-specific binding. **(a)** Mutation enrichment across CBS groups in metastatic tumors. CBS groups were analyzed using RM2 in HMF cancer cohorts with at least 25 samples. Dot color indicates fold change in mutation enrichment relative to flanking regions, and dot size reflects FDR-adjusted significance. Significant increases are highlighted by black outlines (FDR < 0.05). **(b)** Mutation enrichment profiles across CBS groups in colorectal cancer. Mutation density was calculated by aligning CBS at their midpoints. Red regions indicate the core CBS (±25 bps), and grey regions indicate flanking sequences (50 bps). Blue lines show LOESS-smoothed mutation density (span = 0.4) and shading indicates standard error.

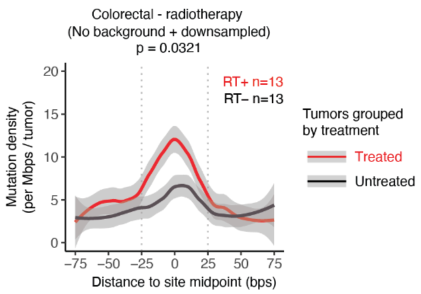


Figure S2. CBS mutation enrichment in radiotherapy-only tumors compared to matched treatment-naïve tumors.

Stringent comparison of tumors treated only with radiotherapy and treatment-naïve tumors. Treatment-naïve tumors were downsampled to match both sample number and rectal cancer proportion of radiotherapy-only tumors. Treatment-naïve tumors were downsampled to match sample size and rectal cancer proportion of radiotherapy-only tumors. Radiotherapy-only tumors exhibited increased CBS mutation enrichment relative to untreated tumors. The y-axis shows aggregated mutation density across all CBS (mutations per Mbps per tumor). Lines show LOESS-smoothed mutation density (span = 0.4) and the shading indicates standard error. Sample sizes and significance of CBS mutation enrichment relative to flanking regions are shown for treated (red text) and untreated groups (black text). The x-axis denotes sequence distance (bps) from CBS midpoints. Gray dotted lines indicate CBS boundaries (50 bps).

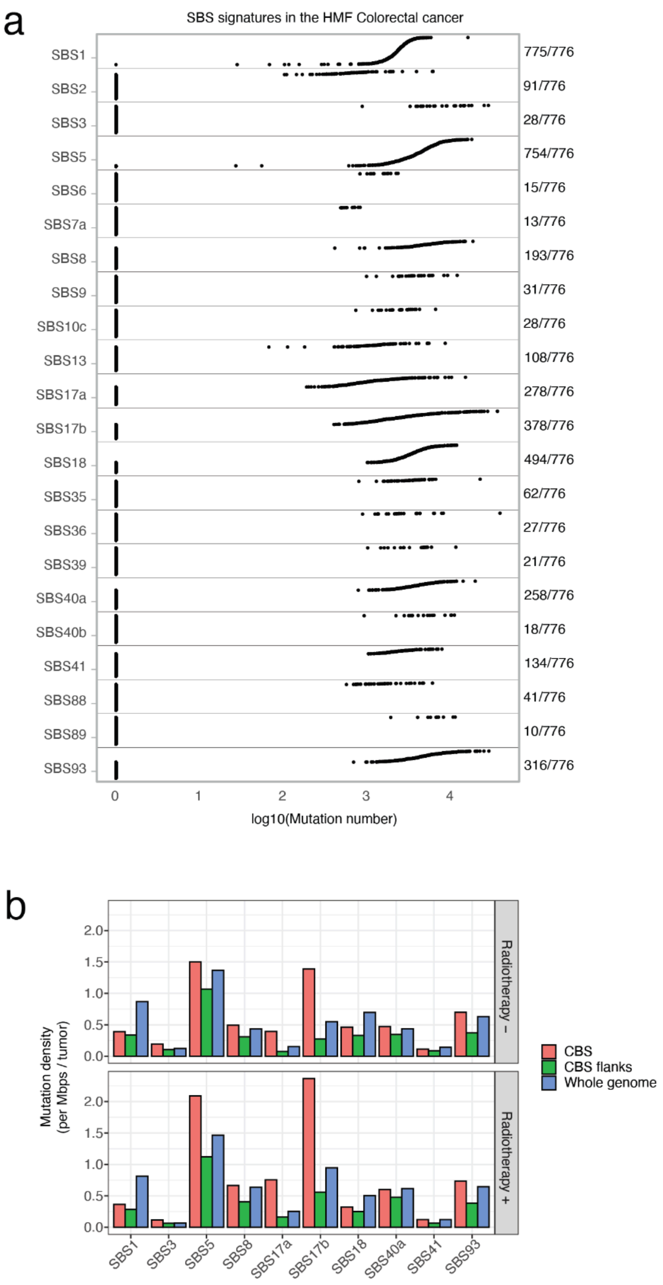


Figure S3. SBS signatures in the colorectal cancer cohort.

**(a)** Prevalence of SBS signatures in metastatic colorectal cancer. Mutation counts reflect assignment solution activity estimated by SigProfilerAssignmentR. Values on the right indicate the fraction of tumors in which each signature was detected. Only signatures present in at least 10 tumors are shown. **(b)** Regional mutation density of SBS signatures at CBS in colorectal tumors grouped by radiotherapy exposure. Only signatures with at least 10 mutations in CBS are shown.

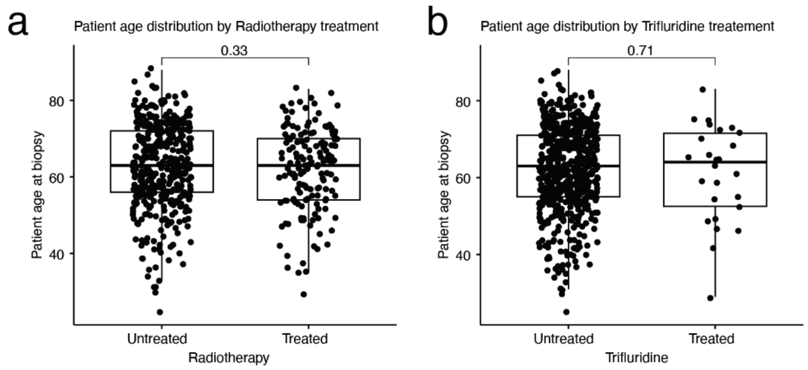


Figure S4. Patient age at biopsy in radiotherapy and trifluridine treatment groups in metastatic colorectal cancer.

Patient age at biopsy compared between **(a)** radiotherapy-treated and untreated tumors and **(b)** trifluridine-treated and untreated tumors. P-values were calculated using two-tailed Wilcoxon tests.

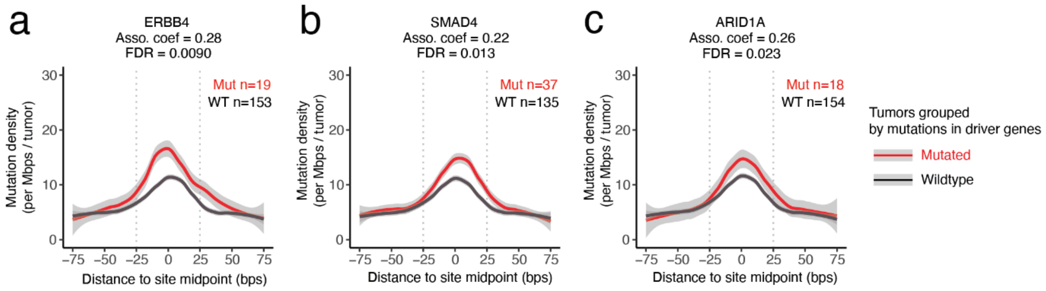


Figure S5. Additional driver gene alterations associated with CBS mutation enrichment in radiotherapy-treated colorectal tumors

Comparison of CBS mutation enrichment in radiotherapy-treated colorectal tumors stratified by driver mutation status. Alterations of *ERBB4*, *SMAD4*, and *ARID1A* were associated with increased CBS mutation enrichment, although effect sizes were modest. The y-axis shows aggregated mutation density across all CBS (mutations per Mb per tumor). Lines show LOESS-smoothed mutation density (span = 0.4) and the shading indicates standard error. Sample sizes and significance of CBS mutation enrichment relative to flanking regions are shown for mutated (red text)and wild-type groups (black text). The x-axis denotes sequence distance (bps) from CBS midpoints. Gray dotted lines indicate CBS boundaries (50 bps).

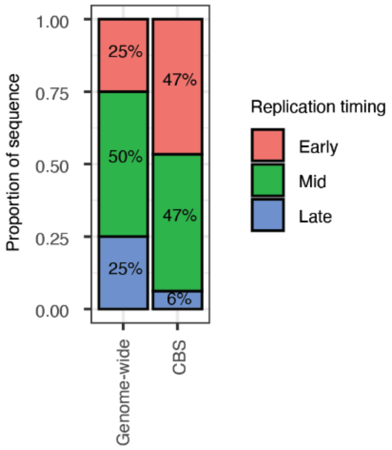


Figure S6. Replication timing distribution of CTCF binding sites.

Stacked bar plots show the distribution of CTCF binding sites (CBS) across replication timing categories compared to the genome-wide background. The left bar represents the genome-wide distribution of replication timing (early 25%, mid 50%, late 25%), and the right bar shows the proportion of CBS in each category. CBS are enriched in early- and mid-replicating regions (47% each) compared to late-replicating regions (6.9%).
